# Cell-type specific autophagy in human leukocytes

**DOI:** 10.1101/2024.12.08.627423

**Authors:** Linh VP Dang, Alexis Martin, Julian M Carosi, Jemima Gore, Sanjna Singh, Timothy J Sargeant

**Affiliations:** Lysosomal Health in Ageing, Lifelong Health, South Australian Health and Medical Research Institute (SAHMRI), Adelaide, SA 5000, Australia; School of Biological Sciences, Faculty of Sciences, Engineering and Technology, The University of Adelaide, Adelaide, SA 5000, Australia; SAHMRI Clinical Trials Platform (CTP), South Australian Health and Medical Research Institute (SAHMRI), Adelaide, SA 5000, Australia; Adelaide Medical School, The University of Adelaide, Adelaide, SA 5000, Australia

**Keywords:** Autophagy, human, autophagic flux, blood, aging, sex

## Abstract

Autophagy is a naturally conserved mechanism crucial for degrading and recycling damaged organelles and proteins to support cell survival. This process slows biological ageing and age-related disease in preclinical models. However, there has been little translation of autophagy to the clinic, and we have identified a lack of measurement tools for physiological human autophagy as a barrier. To address this, we have previously developed a direct measurement tool for autophagy in pooled human peripheral blood mononuclear cells (PBMCs) in the context of whole blood. In order to better understand how autophagy behaves and changes in humans, we measured human autophagic flux using flow cytometry in 19 cell sub-populations in whole blood to retain physiological flux. Autophagic flux was different between different cell types, being highest in B lymphocytes and lowest in T lymphocytes and monocytes. Autophagic flux also varied with sex, being higher in monocytes in females compared with males. In keeping with previous observations in humans, autophagy also increased with ageing at sub-population levels. Importantly, we found that only monocytes – specifically, non-classical monocytes – displayed increased autophagic flux following amino acid withdrawal, underscoring the importance of population selection for measurement of autophagic flux during nutrient restriction studies in humans. Collectively, these data show PBMC population level analysis improves sensitivity of human autophagic flux measurement.

## Introduction

Macroautophagy (hereafter referred to as autophagy) is a nutrient- and stress-responsive process that supports cellular resilience by recycling intracellular material. This allows the provision of nutrients during starvation, clearance of invading pathogens, suppression of inflammation, and maintenance of the mitochondrial network (1–4). The consequences of poorly functioning autophagy in mouse models include accelerated biological aging (4–6), and age-related diseases that include atherosclerosis, dementia, and different cancers (7). Autophagy is also modifiable in cell and preclinical models using pharmacological and nutrition-based interventions (8, 9). This means that autophagy has huge potential as a pathway that could slow or delay the onset of age-related disease.

Although autophagy has immense translational potential, autophagy-based research generally has not progressed beyond preclinical models, although there have been rare exceptions (10–12). The reason for this is in part because autophagy is very difficult to measure in people (13). To address this block to translation, we recently developed a method of measuring autophagic flux in peripheral blood mononuclear cells (PBMCs) by treating whole blood with the lysosome inhibitor chloroquine (CQ). PBMCs are then isolated, and microtubule associated protein 1 light chain 3 beta (MAP1LC3B) isoform II/LC3B-II is measured to derive autophagic flux (14), a dynamic process that analyzes the rate at which autophagy captures and degrades intracellular substrates in the lysosome. Maintaining PBMCs in whole blood during blockade of lysosomal degradation is important because it leaves intact physiological concentrations of nutrients and hormones such as insulin.

Further, in order to develop more precise measures of autophagy in humans that are robust to changes in proportions of cellular populations in the PBMC pool, we need to understand what autophagy looks like in specific cellular sub-populations. This is especially important as development and ageing (15–17), obesity and weight loss (18), exercise (19), and high fat food (20) can all impact cell characteristics and population fractions within the PBMC pool. We aimed to address this issue by developing a physiological autophagic flux measurement using flow cytometry in young and middle-aged adults to evaluate autophagic flux in different cell populations. Understanding these differences will allow for correct interpretation of autophagy measurements in human blood in future clinical trials, and the development of recommendations for targeting PBMC sub-populations in a study-appropriate manner.

## Results

### Physiological autophagic flux retained in whole blood but not in RPMI or plasma

LC3B-II flux was defined as the difference in LC3B-II abundance with the lysosomal inhibitor chloroquine (CQ) minus the LC3B-II abundance in a parallel sample where CQ was not added. While autophagic flux in PBMCs cultured in whole blood or culture media (RPMI) has been successfully measured using western blot and ELISA (12, 21), and LC3B levels have been analyzed in various leukocyte populations using flow cytometry (12, 21–23), physiological autophagic flux in whole blood has not been measured in different cell types using flow cytometry. We, therefore, aimed to do this by culturing whole blood without and with CQ and analysing autophagy in cell types using flow cytometry for LC3B-II. To evaluate antibody specificity for measuring LC3B-II flux, we analysed wildtype and LC3B knockout (KO) HEK 293T cells. We observed a higher signal for LC3B-II in CQ-treated samples compared to control samples, but only in wildtype HEK cells—not in the LC3B KO cells (Figure S1A, B). Similarly, LC3B-II flux was observed in PBMCs (Figure S1C) where LC3B staining was appreciably higher than the IgG isotype control.

We subsequently investigated physiological LC3B-II flux (measured in the context of whole blood) of major blood cell types by flow cytometry including monocytes (classical, intermediate and non-classical monocytes), T lymphocytes (CD4 T lymphocytes: T regulatory cells (Treg), naïve, central memory (CM), effector memory (EM), and terminally differentiated (TEMRA), and CD8 T lymphocytes: naïve, CM, EM and TEMRA), B lymphocytes (naïve and memory B cells), natural killer (NK) cells (CD56^hi^C16^-^, CD56^hi^CD16^+^, CD56^dim^CD16, and CD56^dim^CD16^+^ NK cells) (17, 24) (gating strategy is shown in Figure S1D-V).

LC3B-II flux was measured in different environments including by addition of lysosomal inhibitors to whole blood, nutrient-rich artificial media (RPMI containing 10% FBS, a common culture medium for PBMCs) or in diluted cognate plasma/Dulbecco’s phosphate-buffered saline (DPBS) mixed in a ratio of 1:1 (Figure 1A-F, S2A-J). We observed a significant reduction in overall LC3B-II flux in PBMCs cultured in RPMI containing 10% FBS (Figure 1A-C) or plasma/DPBS (Figure 1D-F) compared to PMBCs exposed to CQ in the context of whole blood. This was particularly evident in NK cells (Figure 1B, E), non-classical monocytes in RPMI medium containing 10% FBS (Figure 1C), and intermediate and non-classical monocytes in cognate plasma/DPBS (Figure 1F).

**Figure 1.**
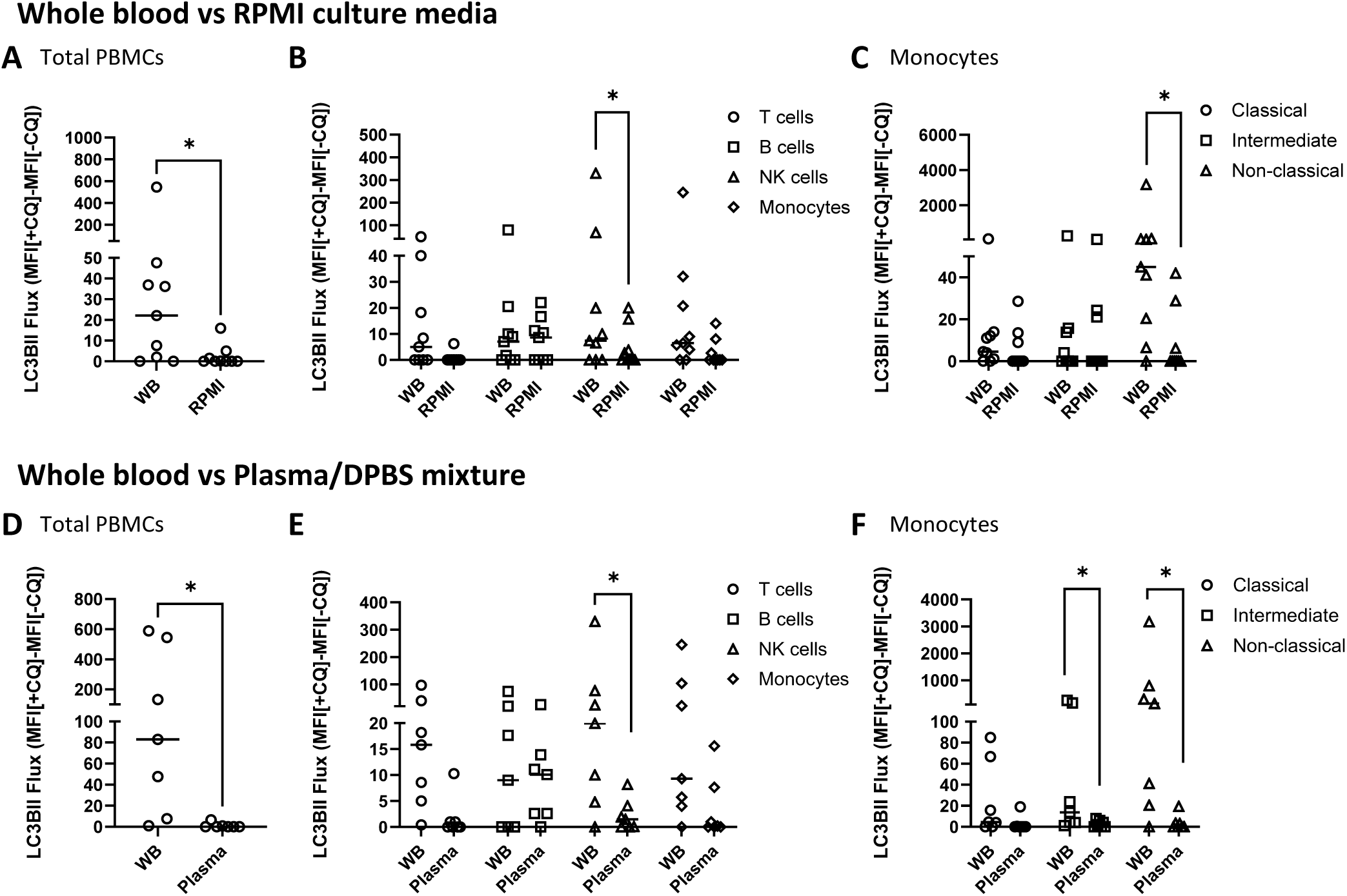
Autophagic flux measurement in physiological and non-physiological environments. LC3B-II flux in PBMCs exposed to CQ in whole blood compared to RPMI containing 10% FBS (N = 9) is shown as follows: Total PBMCs (A) T cells, B cells, NK cells, monocytes (B), Monocyte sub-populations: classical, intermediate, non-classical monocytes (C). LC3B-II flux in PBMCs exposed to CQ in whole blood compared to 1:1 cognate plasma:DPBS (N = 7) is shown as follows: Total LC3B-II flux (D), T cells, B cells, NK cells, monocytes (E), Monocyte sub-populations: classical, intermediate, non-classical monocytes (F). Wilcoxon matched-paired signed rank tests were used. *p<0.05. Bars = median.

### Basal autophagic flux is intrinsically different in different sub-populations of the PBMC pool

Assessing physiological autophagy at the level of leukocyte sub-populations is important to understand how autophagic flux in the PBMC pool as a whole may be impacted by shifting sub-population fractions, which is known to occur with factors such as ageing (17). To understand what cell-type specific physiological autophagic flux looks like, we analysed LC3B-II flux in whole blood in 43 participants. Participant characteristics are presented in Table S1.

Among broad cell type classifications, no significant differences in LC3B-II flux were observed across different cell types (Figure 2A, B). Within each broad cell population, non-classical monocytes had the highest LC3B-II flux of the monocytes (Figure 2C, D); CD56^dim^CD16^+^ cells had the highest flux within the NK cells (Figure 2E); naive B cells exhibited significantly higher LC3B-II flux compared to memory B cells (Figure 2F); and higher LC3B-II flux was found in NKT cells compared to CD4 and CD8 T cells (Figure 2G). Granulocytes showed very high LC3B-II flux (Figure S3A). The proportion of flux contributed by each cell type to the overall flux of the PBMC pool varied significantly from person to person, both in absolute (Figure S3B) and relative amounts (Figure S3C).

**Figure 2.**
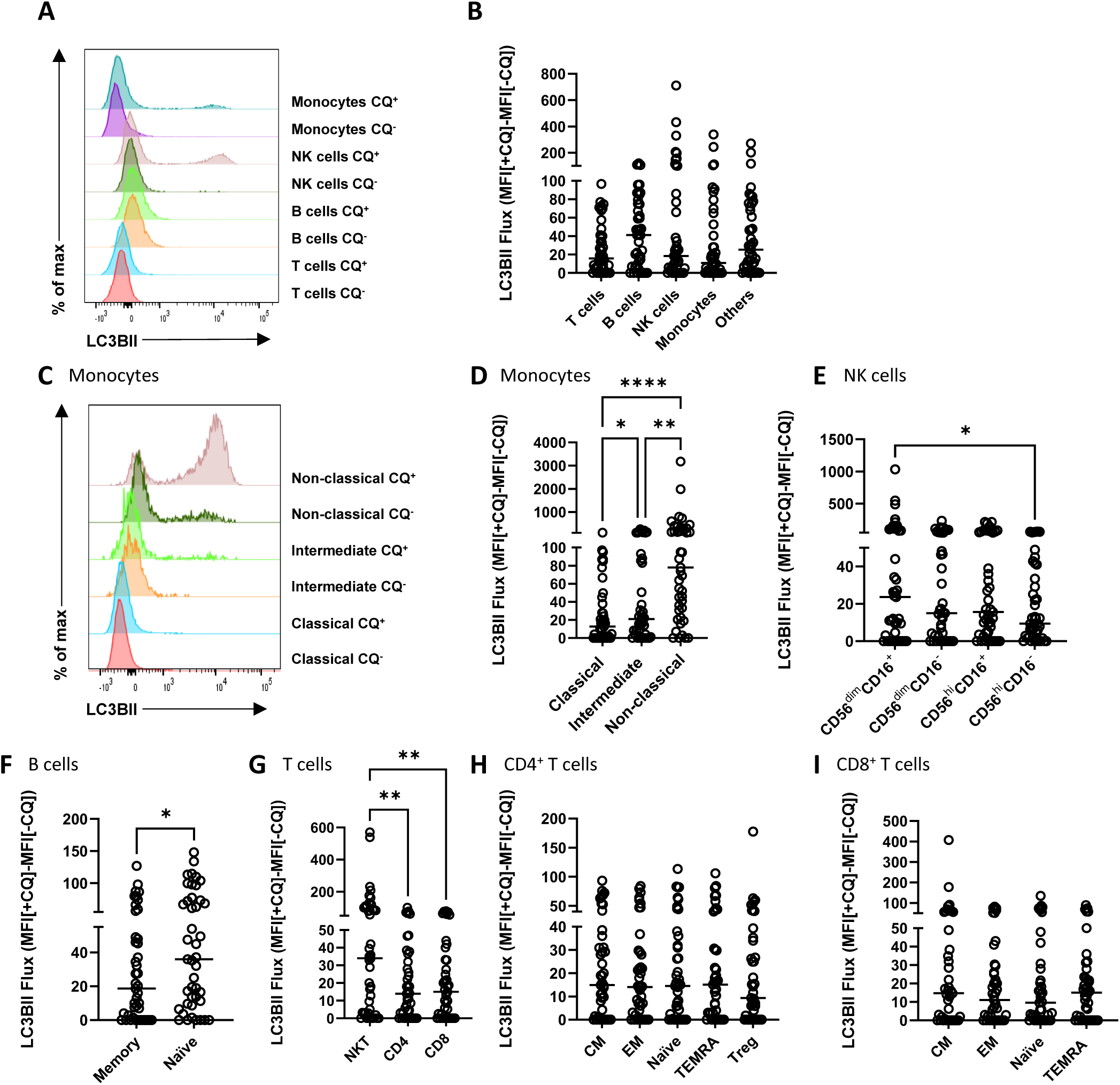
Basal physiological autophagic flux is intrinsically different in different cell types. Flow cytometry histogram representing LC3B-II fluorescent signal in whole blood treated with or without CQ (A). Summary data of LC3B-II flux in different cell populations, including T cells, B cells, NK cells, monocytes, and others (B). Histogram of LC3B-II levels of monocyte sub-populations in whole blood treated with or without CQ including classical, intermediate, and non-classical monocytes (C). Summary data of LC3B-II flux in different sub-populations including Monocyte (D), NK cells (E), B cells (F), T cells, CD4 T cells (H), CD8 T cells (I). Measurement of LC3B-II flux was performed in N = 43 people and presented with median (Friedman test with multiple comparison and Wilcoxon matched-paired signed rank test analysis, *p<0.05, **p<0.01, ***p<0.001).

### Basal autophagic flux in different cell types is highly correlated

While we observed differences in basal LC3B-II flux across various sub-populations, we next investigated whether the flux of each individual cell type correlates with the total PBMC flux. The correlation matrix showed statistically significant correlation between the flux of the total PBMCs and individual cell populations (T cells, B cells, NK cells and monocytes) (Figure 3A and Table S2A) as well as sub-populations (Figure 3B and Table S2H). Correlation was also statistically significant among each cell type (T cells, B cells, NK cells and monocytes) and their corresponding sub-populations (Figure S3D-I, Table S2B-G). The results suggest that basal LC3B-II flux of individual cell populations is highly correlated with the total flux of the whole blood under physiological conditions.

**Figure 3.**
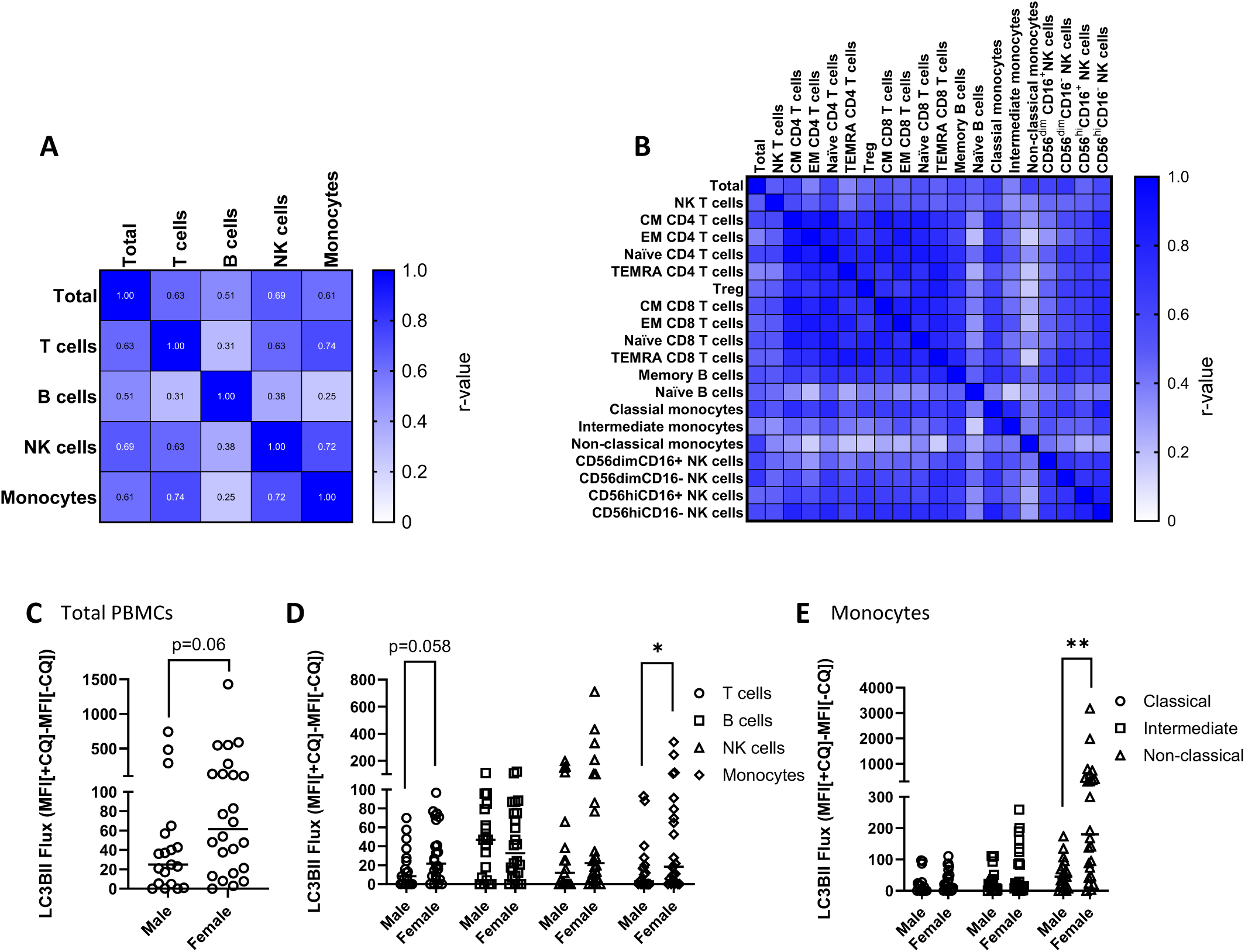
Autophagic flux in different cell types is correlated and shows sex-related differences. A heat map showing Spearman r value for the correlation between total LC3B-II flux and LC3B-II flux of individual cell populations including T cells, B cells, NK cells and monocytes (A) and flux of individual cell sub-populations (B). Graph showing the difference in LC3B-II flux between male and female in total PBMCs (C) and PBMC populations: T cells, B cells, NK cells and monocytes (D) and monocyte sub-populations (E) analysed by Mann-Whitney test, *p<0.05, **p<0.01. Bars = median.

Sex differences in autophagy have been reported in various animal studies (27, 28). We also analyzed the basal LC3B-II flux with regard to sex and observed a trending difference in total LC3B-II flux between males and females (Figure 3C (p = 0.06) and Table S3). However, when examining individual cell populations, we noted that female has significantly higher LC3B-II flux compared to male in monocytes (Figure 3D, Table S3), particularly in non-classical monocytes (Figure 3E, Table S3).

### Basal autophagic flux positively correlates with age

As age has been reported to be positively correlated with autophagic flux in the PBMC pool in older individuals aged 35-70 in people at risk of type-2 diabetes (29), we sought to investigate whether this is also true for healthy younger individuals. Crucially, we also wanted to determine whether PBMC sub-populations also exhibited increases in autophagy with age. We observed a positive correlation between age and LC3B-II flux in total PBMCs (Figure 4A); and other populations including monocytes (Figure 4B), NK cells (Figure 4C); and sub-populations such as T cell subpopulations (including CD8, TEMRA CD8, and TEMRA CD4 T cells, Figure 4D-F)classical monocytes (Figure 4G), intermediate monocytes (Figure 4H), and all NK cell sub-populations (Figure 4I-L). Non-classical monocytes trended with a p-value of 0.053, R^2^=0.09 (data not shown). However, after normalising for sex, autophagic flux of monocytes (p=0.07, R^2^=0.15) and classical monocytes (p=0.89, R^2^=0.15), lost significance. However, other significant correlations remained including total PBMCs (p=0.01, R^2^=0.18), NK cells (p=0.01, R^2^=0.18); and T cell sub-populations (TEMRA CD4 T cells (p=0.02, R^2^=0.14), CD8 T cells (p=0.03, R^2^=0.15), TEMRA CD8 T cells (p=0.03, R^2^=0.11)), intermediate monocytes (p=0.03, R^2^=0.16), NK cell sub-populations (CD56^dim^CD16^+^ NK cells (p=0.02, R^2^=0,15), CD56^dim^CD16^-^ NK cells (p<0.01, R^2^=0.23), CD56^hi^CD16^+^ NK cells (p<0.01, R^2^=0.19), and CD56^hi^CD16^-^ NK cells (p=0.01, R^2^=0.15)).

**Figure 4.**
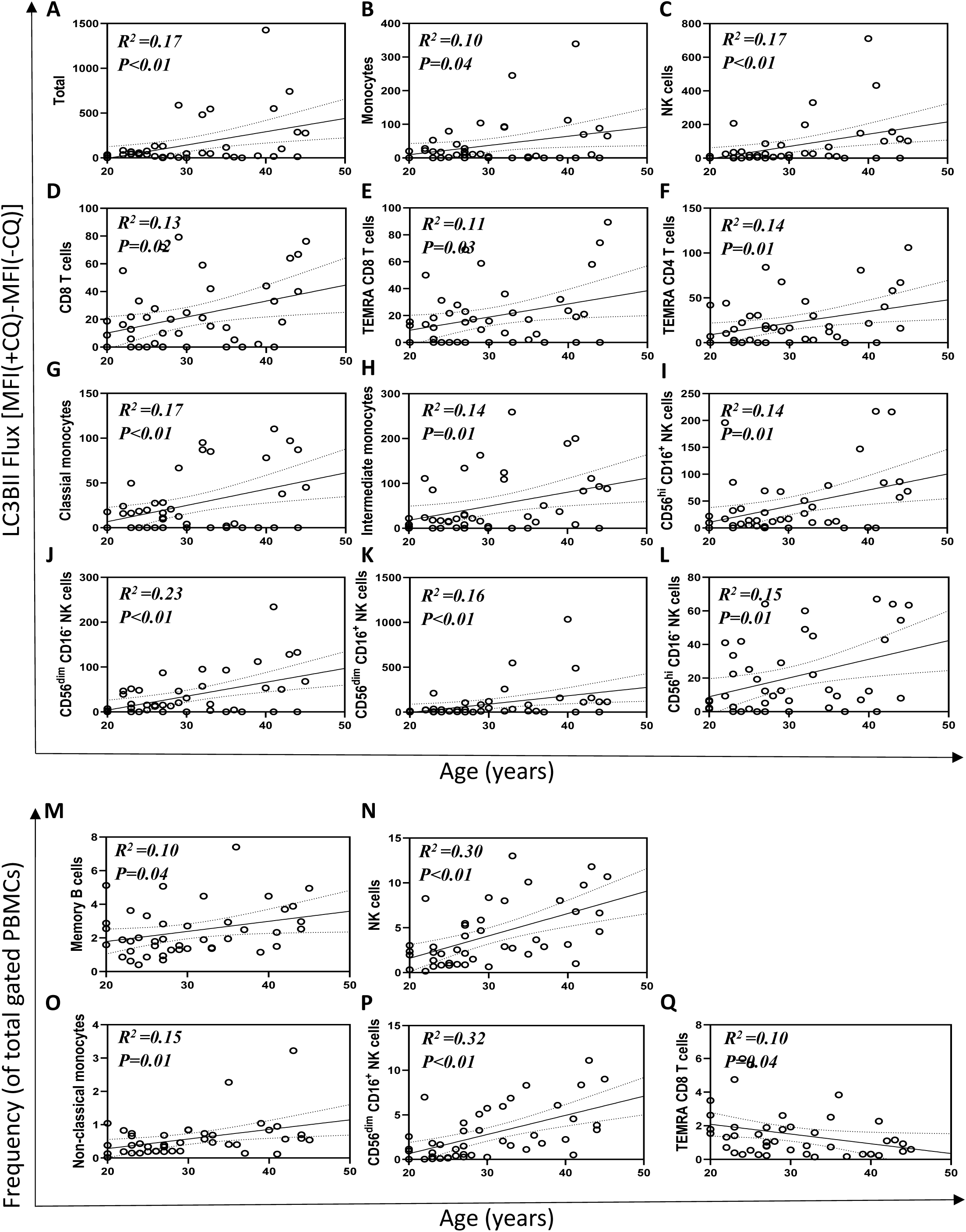
Autophagic flux increases with human age at the cell-type level. Linear regression between age and LC3B-II flux (r^2^ and p value are displayed on graphs) of total PBMCs (A); Monocytes (B); NK cells (C) CD8 T cells (D), TEMRA CD8 T cells (E) and TEMRA CD4 T cells (F); classical monocytes (G), intermediate monocytes (H) CD56^dim^CD16^+^ NK cells (I), CD56^dim^CD16^-^ NK cells (J), CD56^hi^CD16^+^ NK cells (K) and CD56^hi^CD16^-^ NK cells (L). Linear regression between age and proportions of cell populations of total PBMC pool including NK cells (M), memory B cells (N), non-classical monocytes (O), CD56^dim^CD16^+^ NK cells (P) and TEMRA CD8 T cells (Q).

Certain cell populations are reported to change with age with regards to abundance (17), therefore we performed regression analyses and found significant positive correlation of age with frequency of NK cells (Figure 4M); memory B cells (Figure 4N), non-classical monocytes (Figure 4O), and CD56dimCD16+ NK cells (Figure 4P), while TEMRA CD8 T cells showed a negative correlation (Figure 4Q). After normalising for the sex in the analysis we found only NK cells (p<0.01, R^2^=0.31); and TEMRA CD8 T cells (p=0.02, R^2^=0.14), non-classical monocytes (p=0.01, R^2^=0.16), and CD56^dim^CD16^+^ NK cells (p<0.01, R^2^=0.32) retained their significant correlation. Additionally, only percentage of non-classical monocytes (p=0.01, R^2^=0.15) was significantly correlated with total LC3B-II flux and the correlation was even stronger after normalising for age and sex (p<0.01, R^2^=0.82) by linear regression analysis.

### Autophagy induced by nutrient restriction shows cell-type specificity

Autophagic flux is activated by nutrient restriction, because nutrients regulate mTORC1, which inhibits the initiation of autophagy (30). We wanted to determine whether sensitivity to nutrient restriction was cell-type specific as this has implications for how human autophagy is monitored during nutritional intervention studies.

We cultured PBMCs in RPMI formulations with amino acids (aa^+^) and without amino acids (aa^-^), both containing 10% dialyzed fetal bovine serum (dFBS) for 1 h at 37°C and observed no change in total LC3B-II flux (Figure 5A). However, significant increases in LC3B-II flux was observed in monocytes (Figure 5B) and non-classical monocytes (Figure 5C). Glucose did not elicit a strong response (Figure S4A, B) during the incubation period. When data for autophagic flux was categorised into groups of 20-35 or 35-50 years of age, LC3B-II flux of monocytes showed no significant difference between the aa^-^ and aa^+^ conditions (Figure 5D). However, significant higher autophagic flux in response to amino acid restriction was observed in non-classical monocytes in the 35-50 year old group (Figure 5E), but not in the 20-35 year old group. Significantly higher LC3B-II flux was observed only in females for both monocytes (Figure 5F) and non-classical monocytes (Figure 5G) in aa^-^ compared to aa^+^ condition. Lack of significance in males was likely due to a lack of statistical power rather than a lack of effect. Surprisingly, even though total NK cells were not significantly different with regards to amino acid restriction, we observed increased autophagic flux in CD56^dim^CD16^-^ (Figure 5H) and CD56^hi^CD16^+^ NK cells (Figure 5I) as the result of aa withdrawal in females.

**Figure 5.**
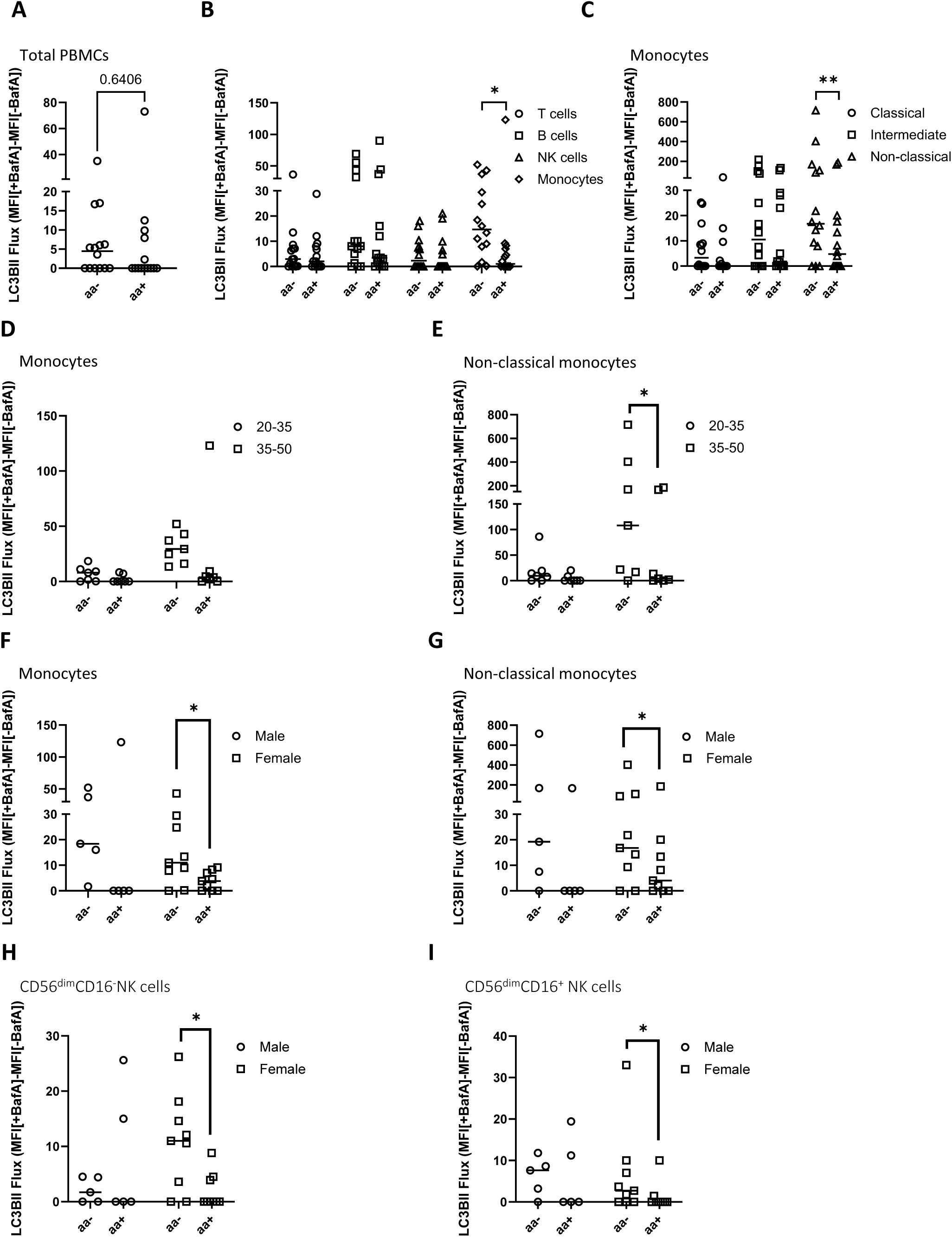
Analysis of LC3B-II flux of isolated PBMCs cultured under amino acid restriction. LC3B-II flux was measured in isolated PBMCs cultured in amino acid-free RPMI containing 10% dFBS (aa^-^) or the same medium spiked with amino acids (aa^+^) with or without BafA (N = 14) as follows: Total PBMCs (A), cell populations: T cells, B cells, NK cells, and monocytes (B) monocyte sub-populations (C). These differences were further stratified into age group (20-35 and 35-50) and presented with LC3B-II flux of monocytes (D) and non-classical monocytes (E). Statistical differences in LC3B-II between aa^-^ and aa^+^ were also analyzed with respect to sex for monocytes (F), non-classical monocytes (G), CD56^dim^CD16^-^ (H) and CD56^hi^CD16^+^NK cells (I). Wilcoxon matched-paired signed rank test was used (*p<0.05, **p<0.01). Bars = median.

## Discussion

In the current study, we confirmed the importance of using whole blood for measuring physiological autophagic flux. We further found that autophagic flux showed cell-type specific variation, and that a positive correlation between age and autophagic flux exists at the total PBMC pool and individual cell type levels. Sex also impacted autophagic flux. Additionally, we identified monocytes, particularly non-classical monocytes, as the most nutrient-sensitive cells, making them well-suited for nutritional intervention studies. Together, these observations provide useful guidance on how to monitor human autophagy in a study appropriate manner.

Consistent with previous research (29), we observed a positive correlation between age and basal autophagic flux. While this had previously been reported in individuals aged 35-75 that were on average obese (29), the current study is the first to report a positive correlation between age and autophagy in a young to middle-aged adult cohort, and importantly at the leukocyte sub-population level. We found a positive correlation between age and autophagic flux in monocytes, NK cells and T lymphocytes and their sub-populations. Other research in human samples showed inconsistent results. While some reported decreased overlap between MAP1LC3B/LC3B and the acidic compartment in CD8 T cells in older individuals (23), and decreased LC3B-II levels in from older compared (31, 32); others showed increased autophagosomes in CD4 T cells in older individuals along with inconsistent changes in autophagy-gene expression (33). However, work from previous relevant studies has not documented autophagic flux and has performed experiments under non-physiological conditions. The relevance of these findings to our work is therefore difficult to ascertain. The data from the current study that show autophagy increases with human ageing in leukocytes suggest that at least in these cell types, autophagy can be upregulated in response to age-related challenges. However, further research is needed to investigate the underlying changes that occur throughout the course of aging that drive this increase in autophagy.

Nutritional withdrawal has been shown to induce autophagy through different mechanisms, including the activation of AMPK and suppression of mTORC1 activity, and subsequent and activation of ULK1, a key initiator of autophagy (reviewed in (34)). In the current study, we observed population specific changes in autophagic flux in response to nutrient restriction. While lymphocyte flux remained relatively stable in different ex-vivo conditions, monocytes (especially non-classical monocytes) exhibited significant responses to amino acid restriction. Consistent with this observation, monocytes are more sensitive than lymphocytes to fasting and exercise with regards to steady-state abundance of autophagy proteins (35); autophagic flux, however, has not previously been assessed. Zhang and colleagues also noted monocytes increase autophagy in response to starvation (36). Although it has been shown that monocytes are nutrient responsive, T lymphocyte autophagy is also likely responsive, albeit to different stimuli not tested in the current study; T lymphocytes increase autophagy in response to growth stimulating factors anti-CD3 antibody and IL-2 (37, 38).

The ability to induce autophagy for nutrient recovery in response to starvation promotes survival under nutrient stress (39, 40). However, this is not the immune system’s only coping mechanism with regards to nutrient restriction. T lymphocytes (41) and monocytes(42, 43) can relocate from the periphery to the nutrient-rich bone marrow during starvation, which has been suggested as a mechanism to protect cells during nutrient restriction. Jordan and colleagues noted the profound migration of Ly-6C^hi^ monocytes (equivalent to classical monocytes in humans) from blood to bone marrow, while the numbers of Ly-6C^lo^ (equivalent to non-classical monocytes in humans) did not differ significantly in mice after four hours of starvation (43). Non-classical monocytes, which can more effectively induce autophagy upon starvation, might survive better in the periphery during nutrient depletion compared to other cell populations that need to migrate immediately to a nutrient-rich environment such as the bone marrow. The changes in cell populations in peripheral blood during starvation suggest the importance of using individual autophagic flux to better monitor changes in autophagy activity. Given their responsiveness to nutrients, monitoring autophagy in monocytes—especially non-classical monocytes—rather than B or T lymphocytes could provide a more sensitive approach to determining whether nutrient restriction promotes autophagy in humans.

The study has several limitations. We used CQ as a lysosomal inhibitor to measure autophagic flux, and this may induce non-canonical autophagy (44). The contribution of this to our measurements is currently unknown. Additionally, we only measured LC3B to probe autophagy, while PBMCs may express other Atg8 proteins, potentially confounding the interpretation. Measuring LC3B-II flux in a cross-sectional study also makes it difficult to conclude that autophagic flux increases with age. Furthermore, selecting participants exclusively from Australia may introduce bias.

Results from this study indicate how best to measure autophagy in human research. We have demonstrated that human cells maintained in artificial media could be subject to large changes in autophagy, meaning that autophagy observed in culture does not resemble autophagy that takes place within a human being. This provides evidence supporting the use of whole blood to assess physiological autophagic flux. We have also shown that autophagy is different in different cellular populations. Thus, changing sub-cellular population fractions within the PBMC pool could impact interpretation of human autophagy studies that measure the whole PBMC pool. We further demonstrated that autophagic flux in humans varies with ageing and sex, even at the cellular sub-population level. This shows that both factors must be considered when designing studies for measurement of autophagy in humans. We also show that autophagy in certain cell types (specifically monocytes/non-classical monocytes) is more sensitive to nutrient restriction than in other cell types – pointing to use of these cells as a better means of measuring autophagy in dietary studies.

## Materials and Methods

Details for reagents and software used in this study can be found in Supplementary Table 4.

### Human ethics approval

The use of the blood samples in the current study was approved by The University of Adelaide Human Research Ethics Committee (Approval HREC H-2021-154). Electronic consent for participation was given after participants had been fully informed of the study.

### Sample collection

Blood samples were collected in lithium heparin Vacuette tubes from study participants between April 2023 and August 2024. The study participants include both men and women aged 20 to 50 years who had normal body weight or were overweight (BMI 18.5-29.9 kg/m2), recruited from Adelaide, with no history of chronic diseases, and without any vaccines or infections within two weeks before the visit. The participants were fasted for a minimum of 12h before blood collection and the samples were processed within 1h of collection. Additional information, including sex, ethnicity, weight, and height, was also collected.

### Treatment of whole blood with CQ

Following collection, 1 mL of whole blood was spilt into two 10 mL conical centrifuge tubes treated with chloroquine diphosphate at a final concentration of 150 µM. The corresponding control tubes were treated with similar amounts of vehicle (water).

### PBMC isolation

PBMCs were isolated using lymphoprep. Blood was mixed with an equal volume of DPBS at a ratio of 1:1 before being carefully layered on top of 15 mL of lymphoprep. PBMCs were collected after being centrifuged for 30 min at 800 x g at room temperature without brake, then washed with DPBS and treated with 1X red blood cell lysis buffer. The PBMCs were then washed twice with DPBS before culturing.

### Culture of isolated PBMCs – incubation in RPMI containing 10% FBS or plasma

In Figures 2 and 3, RPMI containing 10% FBS and diluted cognate plasma (in a 1:1 ratio with DPBS, derived from a lymphoprep-based PBMC isolation, to achieve similar concentration of PBMCs relative to plasma as in whole blood) was directly compared to whole blood as a medium for addition of CQ for lysosomal inhibition. To do this, isolated PBMCs were resuspended in either diluted cognate plasma or RPMI containing 10% FBS and incubated with or without CQ. All samples described here were incubated for 1 h at 37°C with rotation at 10 revolutions per minute using a ThermoFisher Scientific Tube Revolver.

### Culture of HEK 293 T cells

Wild-type and LC3B knockout HEK293T cells were seeded at 0.5 × 10 cells per well in a 12-well plate one day prior to the experiment in Dulbecco’s Modified Eagle’s Medium (DMEM) supplemented with 10% FBS. The cells were treated with CQ or its vehicle for 1 hour at 37°C before performing flow cytometry.

#### CRISPR Cas9 gene editing (MAP1LC3B KO)

Lentiviral vectors (expressing Cas9 and scramble gRNA 5’-GTGTAGTTCGACCATTCGTG or gRNA against human LC3B 5’-CATCCAACCAAAATCCCGGT (pLV[CRISPR]-hCas9:T2A:Bsd-U6>hMAP1LC3B[gRNA#954]) (Vectorbuilder) were transfected into HEK293T cells with plasmids psPAX2 (Addgene #12260) and pCMV-VSV-G (Addgene #8454) using lipofection (Lipofectamine 3000, Thermo Fisher, L3000015). Supernatant containing virus 48 hours post-transfection was passed through a 0.45 µM filter. 8 µg/mL polybrene was added to the filtrate. Target cells (HEK293T) were incubated with virus for 24 h and were left to recover for 48 h before selection (6 µg/mL blasticidin, 7 d). Monoclonal KO cell lines were generated by sorting cells into a 96-well plate at 1 cell/well using a BD FACSAria Fusion. Single cell clones were grown for 2-3 weeks before passaging and analysis. Western blotting was used to verify LC3B KO. psPAX2 was a gift from Didier Trono (Addgene plasmid #12260; http://n2t.net/addgene:12260; RRID:Addgene_12260). pCMV-VSV-G was a gift from Bob Weinberg (Addgene plasmid #8454; http://n2t.net/addgene:8454; RRID:Addgene_8454).

### Culture of isolated PBMCs – nutritional interventions

In Figure 5 and S4, isolated PBMCs were cultured in various nutritional conditions.

Figure 5: amino acid-free RPMI prepared according to manufacturer’s instructions containing 10% dFBS, and amino acid-free RPMI containing 10% dFBS spiked with amino acids, as described in Table S5.

Figure S4 A, B: glucose-free RPMI containing 10% dFBS, and glucose-free RPMI containing 10% dFBS spiked with glucose at a final concentration of 2 g/L.

Duplicate isolated PBMCs were treated with Bafilomycin (BafA) at final concentration of 200 nM and corresponding control tubes were treated with DMSO as a vehicle control. All cultures were incubated at 37°C for 1 h in 12-well plates.

### Flow cytometry staining

Whole blood incubated with CQ and then lysed with red blood cell lysing buffer for 15 minutes at room temperature (RT); PBMCs after being cultured under different conditions; and wild-type and LC3B knockout HEK 293T cells after being cultured with CQ for 1h at 37°C; were harvested, washed twice with DPBS, and incubated for 10 minutes at RT with the recommended amount of fixable viability dye. They were then washed with DPBS before staining with saturated concentrations of surface monoclonal antibodies diluted in BD Horizon Brilliant Stain Buffer Plus for 30 minutes at 4°C (this step was not applied to HEK cells). After staining, cells were washed twice with DPBS and permeabilized with 0.05% saponin for 5 minutes at RT, and then washed with DPBS to remove LC3B-I from the cytosol, leaving only LC3B-II attached to the lysosomal membrane. The cells were then fixed with 4% v/v neutral formalin for 15 mins at RT; washed with DPBS. Finally, samples were incubated with or without anti-LC3B Alexa Fluor-647 (for unstained control) or IgG Alexa Fluor-647 for IgG control (for PBMCs, wildtype, and LC3B knockout HEK 293T cells), diluted 1:50 in DPBS containing 2% bovine serum albumin (BSA) and 0.01% saponin for 1 hour at 4°C. The samples were then washed twice with DPBS containing 2% BSA and 0.01% saponin before flow cytometry analysis using a BD FACSymphony A5 Cell Analyzer (BD Biosciences) at a fixed speed. The FACS Symphony is calibrated daily with BD Cytometer Setup and Tracking Beads.

Surface monoclonal antibodies conjugated to different fluorochromes, including BUV395, BUV615, BUV496, BUV805, BV421, BV480, BV786, BV750, FITC, BB700, PE, PE-CF594, PE-Cy7, Live/dead 780, and Alexa647 were employed in this study to characterize different PBMC sub-populations (detailed in Table S4).

### Quantification and statistical analysis

Flow cytometry data analysis was performed using Flowjo 10.8.0 (BD Biosciences, San Jose, CA, USA) for Windows with a gating strategy performed by a single operator (LVPD) with gating strategy available in Supplementary Figure 1 (D-V). Autophagic flux was quantified as the difference in Geometric Mean Fluorescence Intensity (MFI) of LC3B-II for each cell population in whole blood treated with or without a saturated concentration of CQ of BafA as described by Bensalem et al (21):

ΔMFI (LC3B-II) = (MFI of LC3B-II with CQ) – (MFI of LC3B-II without CQ)
ΔMFI (LC3B-II) = (MFI of LC3B-II with BafA) – (MFI of LC3B-II without BafA).

Graphs were generated using GRAPHPAD PRISM version 10.3.0, for Windows (GraphPad Software, La Jolla, CA, USA) or R version 4.3.2 (R Foundation for Statistical Computing, Vienna, Austria)/RStudio (Integrated Development for R. RStudio, PBC, Boston, MA, USA). Data were analyzed using statistical tests described in figure legends (Friedman test, paired Wilcoxon matched-paired signed rank test, Mann-Whitney test, Spearman correlation and simple linear regression. All tests were performed as two-tailed tests and significant levels are presented as *p<0.05, **p<0.01, or ***p<0.001.

## Acknowledgments.

The authors would like to sincerely thank all the study participants. We thank staff from the SAHMRI Clinical Trial Platform for sample collection, Thanh Thi Dinh, Vuong Tan Phan (Tam Anh Hospital, Hanoi, Vietnam) for assistance with the statistical data analysis, and Dr Randall Grose for flow cytometry support. Flow cytometry analysis was performed at the Adelaide Health and BioMedical Precinct Cytometry Facility (SAHMRI), which is generously supported by the Detmold Group, the McMahon Family, the Australian Cancer Research Foundation, the Cancer Council, and the Australian Government through the Zero Childhood Cancer Program.

## Data availability statement

Raw data supporting the findings of this study are available from the corresponding author on request.

## Author contributions

LVPD and TJS designed all the experiments. LVPD performed all the experiments. LVPD performed all the data analysis. JC designed amino acid supplementation. SS made LC3B KO cells. AM aided in sample collection and processing. JG coordinated the parent human study and human data collection. LVPD and TJS wrote the manuscript. All the authors reviewed the manuscript and provided approval for submission.

## Abbreviations

aa: amino acid
BafA: Bafilomycin A
BSA: Bovine serum albumin
CM: Central memory
CQ: Chloroquine
DPBS: Dulbecco’s phosphate-buffered saline
EM: Effector memory
FBS: Fetal bovine serum
MFI: Geometric Mean Fluorescence Intensity
NK: Natural killer
PBMC: Peripheral blood mononuclear cell
RT: Room temperature
TEMRA: Effector memory cells re-expressing CD45RA
Treg: Regulatory T cell
WB: Whole blood

## References

1. Karsli-Uzunbas, G., Guo, J. Y., Price, S., Teng, X., Laddha, S. V., Khor, S., Kalaany, N. Y., Jacks, T., Chan, C. S., Rabinowitz, J. D., and White, E. (2014) Autophagy is required for glucose homeostasis and lung tumor maintenance. Cancer discovery 4, 914–927

2. Jassey, A., and Jackson, W. T. (2023) Viruses and autophagy: bend, but don’t break. Nat Rev Microbiol

3. Lazarou, M., Sliter, D. A., Kane, L. A., Sarraf, S. A., Wang, C., Burman, J. L., Sideris, D. P., Fogel, A. I., and Youle, R. J. (2015) The ubiquitin kinase PINK1 recruits autophagy receptors to induce mitophagy. Nature 524, 309–314

4. Cassidy, L. D., Young, A. R. J., Young, C. N. J., Soilleux, E. J., Fielder, E., Weigand, B. M., Lagnado, A., Brais, R., Ktistakis, N. T., Wiggins, K. A., Pyrillou, K., Clarke, M. C. H., Jurk, D., Passos, J. F., and Narita, M. (2020) Temporal inhibition of autophagy reveals segmental reversal of ageing with increased cancer risk. Nature Communications 11, 307

5. Carosi, J. M., Fourrier, C., Bensalem, J., and Sargeant, T. J. (2021) The mTOR-lysosome axis at the centre of ageing. FEBS Open Bio

6. López-Otín, C., Blasco, M. A., Partridge, L., Serrano, M., and Kroemer, G. (2023) Hallmarks of aging: An expanding universe. Cell 186, 243–278

7. Klionsky, D. J., Petroni, G., Amaravadi, R. K., Baehrecke, E. H., Ballabio, A., Boya, P., Bravo-San Pedro, J. M., Cadwell, K., Cecconi, F., Choi, A. M. K., Choi, M. E., Chu, C. T., Codogno, P., Colombo, M. I., Cuervo, A. M., Deretic, V., Dikic, I., Elazar, Z., Eskelinen, E. L., Fimia, G. M., Gewirtz, D. A., Green, D. R., Hansen, M., Jäättelä, M., Johansen, T., Juhász, G., Karantza, V., Kraft, C., Kroemer, G., Ktistakis, N. T., Kumar, S., Lopez-Otin, C., Macleod, K. F., Madeo, F., Martinez, J., Meléndez, A., Mizushima, N., Münz, C., Penninger, J. M., Perera, R. M., Piacentini, M., Reggiori, F., Rubinsztein, D. C., Ryan, K. M., Sadoshima, J., Santambrogio, L., Scorrano, L., Simon, H. U., Simon, A. K., Simonsen, A., Stolz, A., Tavernarakis, N., Tooze, S. A., Yoshimori, T., Yuan, J., Yue, Z., Zhong, Q., Galluzzi, L., and Pietrocola, F. (2021) Autophagy in major human diseases. The EMBO journal, e108863

8. Bensalem, J., Fourrier, C., Hein, L. K., Hassiotis, S., Proud, C. G., and Sargeant, T. J. (2021) Inhibiting mTOR activity using AZD2014 increases autophagy in the mouse cerebral cortex. Neuropharmacology, 108541

9. Hein, L. K., Apaja, P. M., Hattersley, K., Grose, R. H., Xie, J., Proud, C. G., and Sargeant, T. J. (2017) A novel fluorescent probe reveals starvation controls the commitment of amyloid precursor protein to the lysosome. Biochim Biophys Acta Mol Cell Res 1864, 1554–1565

10. Rangwala, R., Chang, Y. C., Hu, J., Algazy, K. M., Evans, T. L., Fecher, L. A., Schuchter, L. M., Torigian, D. A., Panosian, J. T., Troxel, A. B., Tan, K. S., Heitjan, D. F., DeMichele, A. M., Vaughn, D. J., Redlinger, M., Alavi, A., Kaiser, J., Pontiggia, L., Davis, L. E., O’Dwyer, P. J., and Amaravadi, R. K. (2014) Combined MTOR and autophagy inhibition: phase I trial of hydroxychloroquine and temsirolimus in patients with advanced solid tumors and melanoma. Autophagy 10, 1391–1402

11. Haas, N. B., Appleman, L. J., Stein, M., Redlinger, M., Wilks, M., Xu, X., Onorati, A., Kalavacharla, A., Kim, T., Zhen, C. J., Kadri, S., Segal, J. P., Gimotty, P. A., Davis, L. E., and Amaravadi, R. K. (2019) Autophagy Inhibition to Augment mTOR Inhibition: a Phase I/II Trial of Everolimus and Hydroxychloroquine in Patients with Previously Treated Renal Cell Carcinoma. Clin Cancer Res 25, 2080–2087

12. Bensalem, J., Teong, X. T., Hattersley, K. J., Hein, L. K., Fourrier, C., Liu, K., Hutchison, A. T., Heilbronn, L. K., and Sargeant, T. J. (2023) Basal autophagic flux measured in blood correlates positively with age in adults at increased risk of type 2 diabetes. Geroscience

13. Sargeant, T. J., and Bensalem, J. (2021) Human autophagy measurement: an underappreciated barrier to translation. Trends Mol Med

14. Bensalem, J., Hattersley, K. J., Hein, L. K., Teong, X. T., Carosi, J. M., Hassiotis, S., Grose, R. H., Fourrier, C., Heilbronn, L. K., and Sargeant, T. J. (2020) Measurement of autophagic flux in humans: an optimized method for blood samples. Autophagy, 1–18

15. Patel, A. A., and Yona, S. (2019) Inherited and Environmental Factors Influence Human Monocyte Heterogeneity. Front Immunol 10, 2581

16. Schefold, J. C., Porz, L., Uebe, B., Poehlmann, H., von Haehling, S., Jung, A., Unterwalder, N., and Meisel, C. (2015) Diminished HLA-DR expression on monocyte and dendritic cell subsets indicating impairment of cellular immunity in pre-term neonates: a prospective observational analysis. J Perinat Med 43, 609–618

17. Kverneland, A. H., Streitz, M., Geissler, E., Hutchinson, J., Vogt, K., Boes, D., Niemann, N., Pedersen, A. E., Schlickeiser, S., and Sawitzki, B. (2016) Age and gender leucocytes variances and references values generated using the standardized ONE-Study protocol. Cytometry A 89, 543–564

18. Poitou, C., Dalmas, E., Renovato, M., Benhamo, V., Hajduch, F., Abdennour, M., Kahn, J. F., Veyrie, N., Rizkalla, S., Fridman, W. H., Sautès-Fridman, C., Clément, K., and Cremer, I. (2011) CD14dimCD16+ and CD14+CD16+ monocytes in obesity and during weight loss: relationships with fat mass and subclinical atherosclerosis. Arterioscler Thromb Vasc Biol 31, 2322–2330

19. Duggal, N. A., Pollock, R. D., Lazarus, N. R., Harridge, S., and Lord, J. M. (2018) Major features of immunesenescence, including reduced thymic output, are ameliorated by high levels of physical activity in adulthood. Aging cell 17

20. Khan, I. M., Pokharel, Y., Dadu, R. T., Lewis, D. E., Hoogeveen, R. C., Wu, H., and Ballantyne, C. M. (2016) Postprandial Monocyte Activation in Individuals With Metabolic Syndrome. J Clin Endocrinol Metab 101, 4195–4204

21. Bensalem, J., Hattersley, K. J., Hein, L. K., Teong, X. T., Carosi, J. M., Hassiotis, S., Grose, R. H., Fourrier, C., Heilbronn, L. K., and Sargeant, T. J. (2021) Measurement of autophagic flux in humans: an optimized method for blood samples. Autophagy 17, 3238–3255

22. Alonzi, T., Petruccioli, E., Vanini, V., Fimia, G. M., and Goletti, D. (2019) Optimization of the autophagy measurement in a human cell line and primary cells by flow cytometry. Eur J Histochem 63

23. Phadwal, K., Alegre-Abarrategui, J., Watson, A. S., Pike, L., Anbalagan, S., Hammond, E. M., Wade-Martins, R., McMichael, A., Klenerman, P., and Simon, A. K. (2012) A novel method for autophagy detection in primary cells: impaired levels of macroautophagy in immunosenescent T cells. Autophagy 8, 677–689

24. Amand, M., Iserentant, G., Poli, A., Sleiman, M., Fievez, V., Sanchez, I. P., Sauvageot, N., Michel, T., Aouali, N., Janji, B., Trujillo-Vargas, C. M., Seguin-Devaux, C., and Zimmer, J. (2017) Human CD56(dim)CD16(dim) Cells As an Individualized Natural Killer Cell Subset. Front Immunol 8, 699

25. Monaco, G., Lee, B., Xu, W., Mustafah, S., Hwang, Y. Y., Carre, C., Burdin, N., Visan, L., Ceccarelli, M., Poidinger, M., Zippelius, A., Pedro de Magalhaes, J., and Larbi, A. (2019) RNA-Seq Signatures Normalized by mRNA Abundance Allow Absolute Deconvolution of Human Immune Cell Types. Cell Rep 26, 1627–1640 e1627

26. Bordi, M., De Cegli, R., Testa, B., Nixon, R. A., Ballabio, A., and Cecconi, F. (2021) A gene toolbox for monitoring autophagy transcription. Cell Death Dis 12, 1044

27. Demarest, T. G., Waite, E. L., Kristian, T., Puche, A. C., Waddell, J., McKenna, M. C., and Fiskum, G. (2016) Sex-dependent mitophagy and neuronal death following rat neonatal hypoxia-ischemia. Neuroscience 335, 103–113

28. Fourrier, C., Bryksin, V., Hattersley, K., Hein, L. K., Bensalem, J., and Sargeant, T. J. (2021) Comparison of chloroquine-like molecules for lysosomal inhibition and measurement of autophagic flux in the brain. Biochem Biophys Res Commun 534, 107–113

29. Bensalem, J., Teong, X. T., Hattersley, K. J., Hein, L. K., Fourrier, C., Liu, K., Hutchison, A. T., Heilbronn, L. K., and Sargeant, T. J. (2023) Basal autophagic flux measured in blood correlates positively with age in adults at increased risk of type 2 diabetes. Geroscience 45, 3549–3560

30. Kim, J., Kundu, M., Viollet, B., and Guan, K. L. (2011) AMPK and mTOR regulate autophagy through direct phosphorylation of Ulk1. Nat Cell Biol 13, 132–141

31. McCormick, J. J., King, K. E., Cote, M. D., Meade, R. D., Akerman, A. P., and Kenny, G. P. (2021) Impaired autophagy following ex vivo heating at physiologically relevant temperatures in peripheral blood mononuclear cells from elderly adults. J Therm Biol 95, 102790

32. Bharath, L. P., Agrawal, M., McCambridge, G., Nicholas, D. A., Hasturk, H., Liu, J., Jiang, K., Liu, R., Guo, Z., Deeney, J., Apovian, C. M., Snyder-Cappione, J., Hawk, G. S., Fleeman, R. M., Pihl, R. M. F., Thompson, K., Belkina, A. C., Cui, L., Proctor, E. A., Kern, P. A., and Nikolajczyk, B. S. (2020) Metformin Enhances Autophagy and Normalizes Mitochondrial Function to Alleviate Aging-Associated Inflammation. Cell Metab 32, 44–55 e46

33. Bektas, A., Schurman, S. H., Gonzalez-Freire, M., Dunn, C. A., Singh, A. K., Macian, F., Cuervo, A. M., Sen, R., and Ferrucci, L. (2019) Age-associated changes in human CD4(+) T cells point to mitochondrial dysfunction consequent to impaired autophagy. Aging (Albany NY) 11, 9234–9263

34. Russell, R. C., Yuan, H. X., and Guan, K. L. (2014) Autophagy regulation by nutrient signaling. Cell Res 24, 42–57

35. Kropfl, J. M., Morandi, C., Gasser, B. A., Schoch, R., Schmidt-Trucksass, A., and Brink, M. (2022) Lymphocytes are less sensitive to autophagy than monocytes during fasting and exercise conditions. Apoptosis 27, 730–739

36. Zhang, Y., Morgan, M. J., Chen, K., Choksi, S., and Liu, Z. G. (2012) Induction of autophagy is essential for monocyte-macrophage differentiation. Blood 119, 2895–2905

37. Li, C., Capan, E., Zhao, Y., Zhao, J., Stolz, D., Watkins, S. C., Jin, S., and Lu, B. (2006) Autophagy is induced in CD4+ T cells and important for the growth factor-withdrawal cell death. J Immunol 177, 5163–5168

38. Alsaleh, G., Panse, I., Swadling, L., Zhang, H., Richter, F. C., Meyer, A., Lord, J., Barnes, E., Klenerman, P., Green, C., and Simon, A. K. (2020) Autophagy in T cells from aged donors is maintained by spermidine and correlates with function and vaccine responses. Elife 9

39. Kuma, A., Hatano, M., Matsui, M., Yamamoto, A., Nakaya, H., Yoshimori, T., Ohsumi, Y., Tokuhisa, T., and Mizushima, N. (2004) The role of autophagy during the early neonatal starvation period. Nature 432, 1032–1036

40. Komatsu, M., Waguri, S., Ueno, T., Iwata, J., Murata, S., Tanida, I., Ezaki, J., Mizushima, N., Ohsumi, Y., Uchiyama, Y., Kominami, E., Tanaka, K., and Chiba, T. (2005) Impairment of starvation-induced and constitutive autophagy in Atg7-deficient mice. J Cell Biol 169, 425–434

41. Collins, N., Han, S. J., Enamorado, M., Link, V. M., Huang, B., Moseman, E. A., Kishton, R. J., Shannon, J. P., Dixit, D., Schwab, S. R., Hickman, H. D., Restifo, N. P., McGavern, D. B., Schwartzberg, P. L., and Belkaid, Y. (2019) The Bone Marrow Protects and Optimizes Immunological Memory during Dietary Restriction. Cell 178, 1088–1101 e1015

42. Janssen, H., Kahles, F., Liu, D., Downey, J., Koekkoek, L. L., Roudko, V., D’Souza, D., McAlpine, C. S., Halle, L., Poller, W. C., Chan, C. T., He, S., Mindur, J. E., Kiss, M. G., Singh, S., Anzai, A., Iwamoto, Y., Kohler, R. H., Chetal, K., Sadreyev, R. I., Weissleder, R., Kim-Schulze, S., Merad, M., Nahrendorf, M., and Swirski, F. K. (2023) Monocytes re-enter the bone marrow during fasting and alter the host response to infection. Immunity 56, 783–796 e787

43. Jordan, S., Tung, N., Casanova-Acebes, M., Chang, C., Cantoni, C., Zhang, D., Wirtz, T. H., Naik, S., Rose, S. A., Brocker, C. N., Gainullina, A., Hornburg, D., Horng, S., Maier, B. B., Cravedi, P., LeRoith, D., Gonzalez, F. J., Meissner, F., Ochando, J., Rahman, A., Chipuk, J. E., Artyomov, M. N., Frenette, P. S., Piccio, L., Berres, M. L., Gallagher, E. J., and Merad, M. (2019) Dietary Intake Regulates the Circulating Inflammatory Monocyte Pool. Cell 178, 1102–1114 e1117

44. Florey, O., Gammoh, N., Kim, S. E., Jiang, X., and Overholtzer, M. (2015) V-ATPase and osmotic imbalances activate endolysosomal LC3 lipidation. Autophagy 11, 88–99

