## Supplementary figures and images for "Cell-type specific autophagy in human leukocytes"

### Figure S

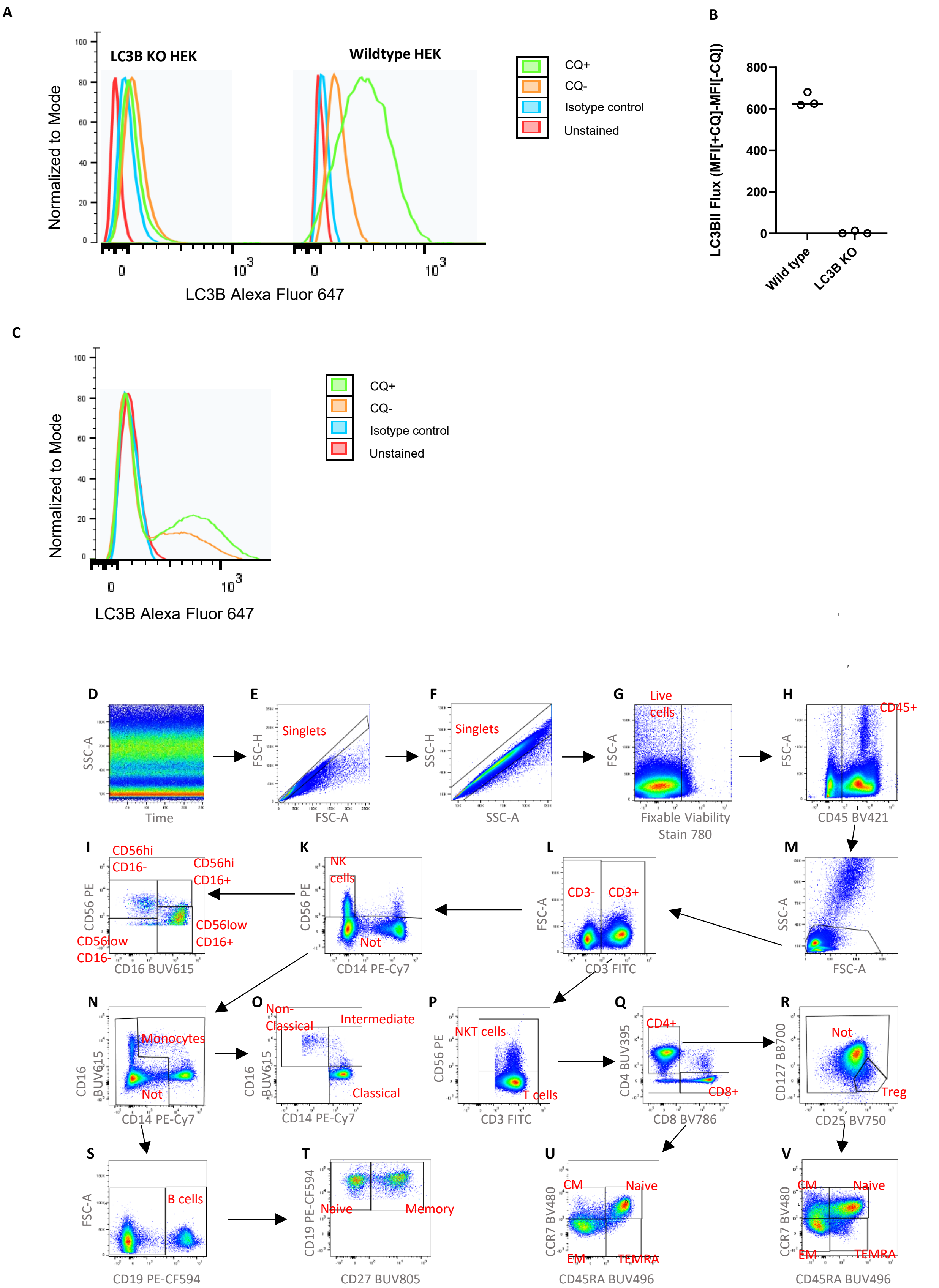

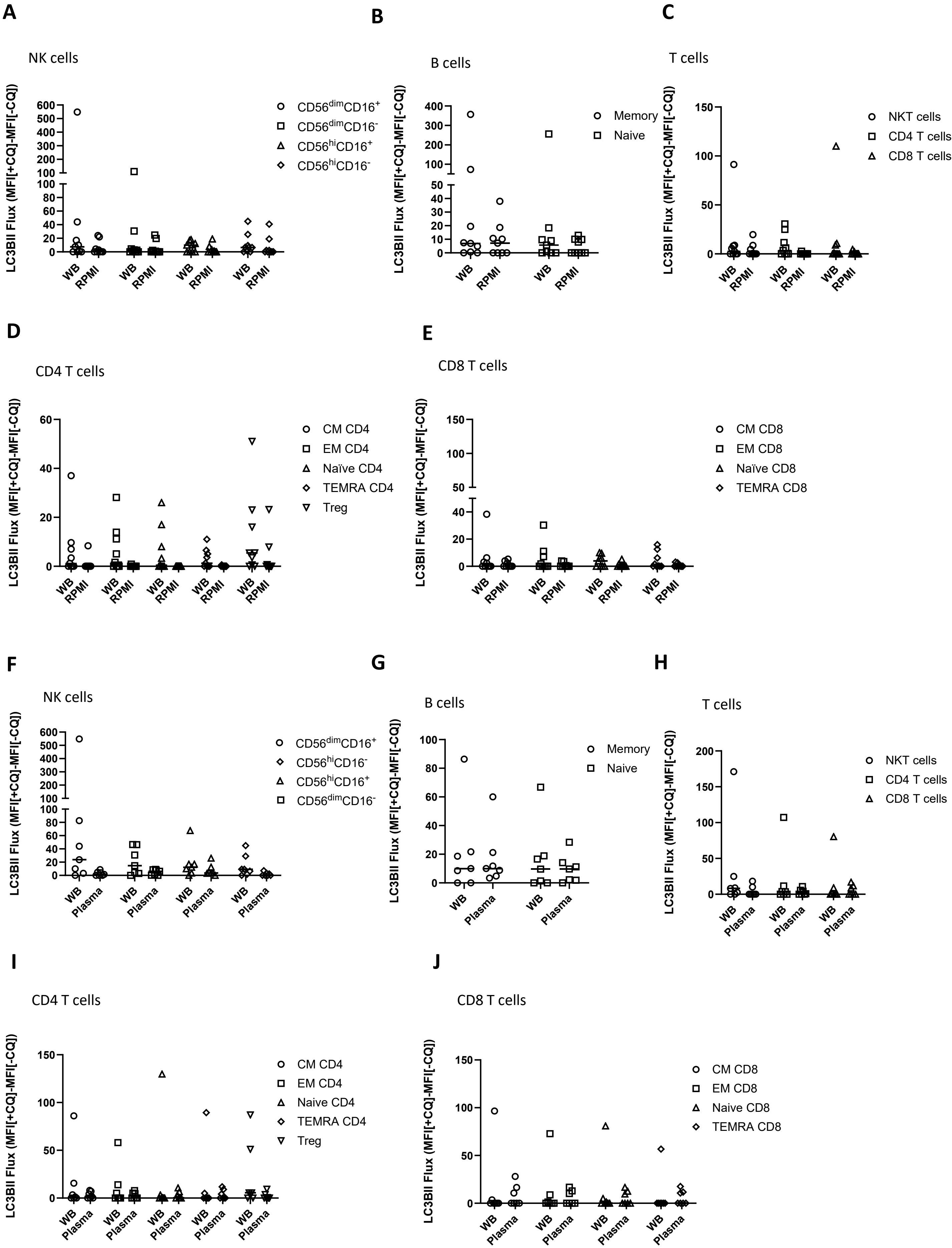

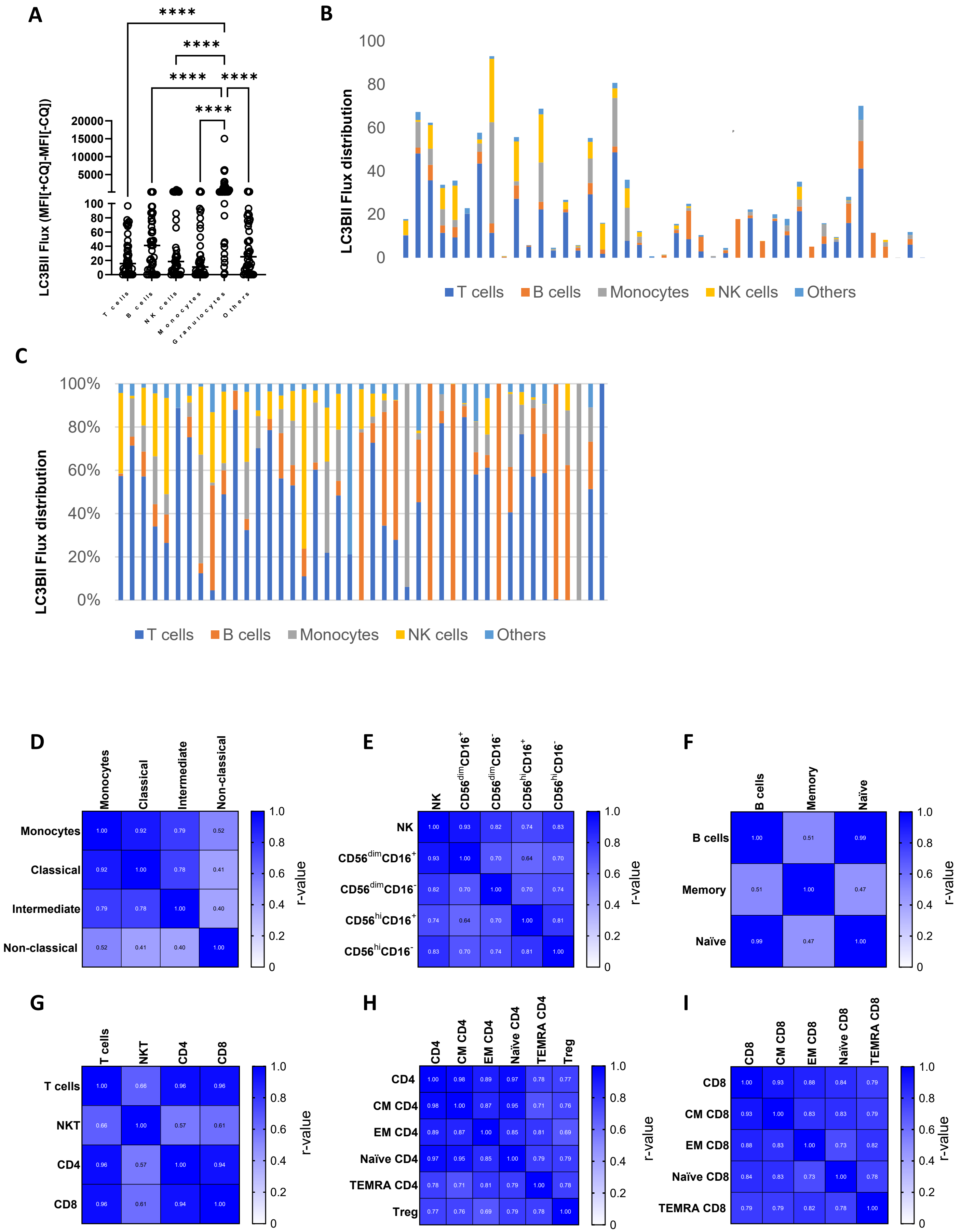

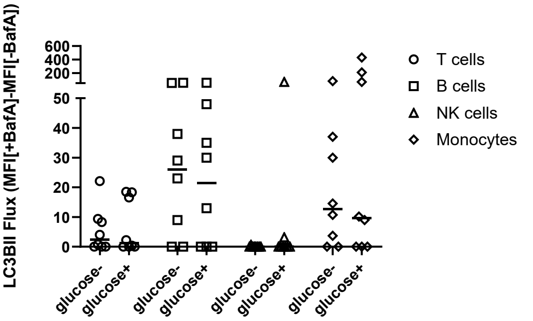

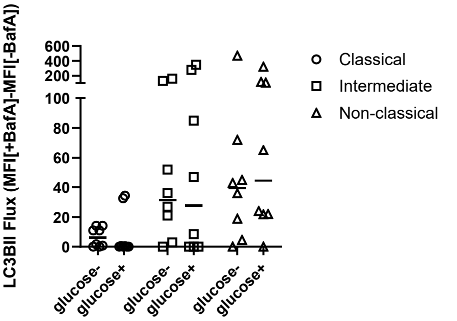
