## Supplementary material for "Cell-type specific autophagy in human leukocytes": Figure S

### Supplementary materials

### Figure legends

***Figure S1.***

Flow cytometry staining of wild type and LC3B knockout HEK 293T cells, treated with or without CQ, with unstained and IgG Alexa 647 conditions as controls (A). The plot represents LC3B-II flux from three independent experiments (B). A histogram represents LC3B-II flux in total PBMCs (C).

Gating strategy for the identification of leukocyte populations in participant whole blood samples by flow cytometry: (D-V)

1. Leukocytes were identified by gating on time (D), single cells (E, F), live cells based on low fluorescence of BD Horizon Fixable Viability Stain 780 (G), selection of $CD45^{+}$ cells (H), followed by FSC and SSC gate for monocytes and lymphocytes (M).
2. T cells were identified by staining for CD3 (L), which were further categorized into Natural Killer T (NKT) cells ($CD3^{+}CD56^{+}$) and other conventional T cells ($CD3^{+}CD56^{-}$) (P). Other conventional T cells were stratified into CD4 and CD8 T cells (Q). CD4 T cells ($CD3^{+}CD4^{+}$) were then classified into the following sub-populations: Treg ($CD25^{+}CD127^{low/-}$) (R) and subsequently divided into naïve ($CD45RA^{+}CCR7^{+}$), central memory ($CD45RA^{-}CCR7^{+}$), effector memory ($CD45RA^{-}CCR7^{+}$), terminally differentiated effector memory (TEMRA; $CD45RA^{+}CCR7^{-}$) (V). A similar gating strategy was applied to CD8 T cell sub-populations, including naïve ($CD45RA^{+}CCR7^{+}$), central memory ($CD45RA^{-}CCR7^{+}$), effector memory ($CD45RA^{-}CCR7^{+}$), TEMRA ($CD45RA^{+}CCR7^{-}$) (U) based on CD45RA and CCR7 expression.
3. NK cells were identified by gating for $CD3^{-}CD56^{+}$ (K), which were then further categorized into $CD56^{hi}CD16^{-}$, $CD56^{hi}CD16^{+}$, $CD56^{dim}CD16^{-}$ and $CD56^{dim}CD16^{+}$ sub-populations (I).
4. Monocytes were gated based on $CD3^{-}CD56^{-}CD14^{+}$ (N), which were then divided into classical monocytes ($CD14^{+}CD16^{-}$), intermediate monocytes ($CD14^{+}CD16^{+}$), and non-classical monocytes ($CD14^{low}CD16^{+}$) (O).
5. B cells were identified by $CD3^{-}CD56^{-}CD14^{-}CD19^{+}$ (S), which were subsequently characterized into naïve ($CD19^{+}CD27^{-}$) and memory B cells ($CD19^{+}CD27^{+}$) (T).

***Figure S2***. Comparison of LC3B-II flux in different populations in whole blood (WB) and isolated PBMCs cultured in RPMI medium containing 10% FBS and presented as subpopulations of the following: NK cells (A), B cells (B), T cells (C), CD4 T cells (D), CD8 T cells (E). LC3B-II flux in different populations in whole blood and isolated PBMC cultured in diluted cognate plasma:DPBS (1:1) and presented as subpopulations of the following: NK cells (F), B cells (G), T cells (H), CD4 T cells (I), CD8 T cells (J).

***Figure S3.*** Plot representing LC3B-II flux of different cell populations including T cells, B cells, NK cells, monocytes, granulocytes and others (A). Distribution of LC3B-II flux of each population contributing to the total LC3B-II flux of the PBMC pool (excluding granulocytes): absolute number (B) and relative percentage (C). Heatmap representing the correlation matrix of individual cell populations and its corresponding subpopulations, including monocytes (D), NK cells (E), B cells (F), T cells (G), CD4 T cells (H), and CD8 T cells (I).

***Figure S4.*** Analysis of LC3B-II flux in different cell types in PBMCs cultured with glucose-free RPMI containing 10% dFCS (glucose-) or the same medium spiked with glucose (glucose+) (N = 8) and presented as follows: T cells, B cells, NK cells, and monocytes (A), and monocyte sub-populations (B).

### Tables

| | | Total (n=43) | Males (n=19) | Female (n=24) | p value |
|---|---|---|---|---|---|
| ***Age (years)*** | | 28 (20-45) | 27(20-44) | 29.5 (20-45) | 0.19 |
| ***Gender, female n (%)*** | | 24 (55.8%) | | | |
| ***Body mass index (BMI, kg/m²)*** | | 23.8 (19.1-30.9) | 25.1 (20.4-30.9) | 21.9 (19.1-29.5) | 0.05 |
| ***Ethnicity*** | | | | | |
| | Caucasian | 18(41.6%) | 6 (31.6%) | 12(50%) | 0.14 |
| | Asian | 24 (58.1%) | 13 (68.4%) | 11(45.8%) | |
| | Others | 1(2.3%) | 0 | 1(4.2%) | |

***Table S1.*** Participant characteristics, analyzed by Mann-Whitney and Kruskal Wallis test.

| **r/p value** | ***Total*** | ***T cells*** | ***B cells*** | ***Monocytes*** | ***NK cells*** |
|---|---|---|---|---|---|
| ***Total*** | | **<0.01** | **<0.01** | **<0.01** | **<0.01** |
| ***T cells*** | 0.63 | | **0.05** | **<0.01** | **<0.01** |
| ***B cells*** | 0.51 | 0.31 | | 0.10 | **0.01** |
| ***Monocytes*** | 0.61 | 0.74 | 0.25 | | **<0.01** |
| ***NK cells*** | 0.69 | 0.63 | 0.38 | 0.72 | |

Table S2A.

| ***r/p value*** | ***T cells*** | ***NKT*** | ***CD4*** | ***CD8*** |
|---|---|---|---|---|
| ***T cells*** | | **<0.01** | **<0.01** | **<0.01** |
| ***NKT*** | 0.66 | | **<0.01** | **<0.01** |
| ***CD4*** | 0.96 | 0.57 | | **<0.01** |
| ***CD8*** | 0.96 | 0.61 | 0.94 | |

Table S2B.

| ***r/p value*** | ***CD4*** | ***CM*** | ***EM*** | ***Naïve*** | ***TEMRA*** | ***Treg*** |
|---|---|---|---|---|---|---|
| ***CD4*** | | **<0.01** | **<0.01** | **<0.01** | **<0.01** | **<0.01** |
| ***CM*** | 0.976 | | **<0.01** | **<0.01** | **<0.01** | **<0.01** |
| ***EM*** | 0.894 | 0.871 | | **<0.01** | **<0.01** | **<0.01** |
| ***Naïve*** | 0.967 | 0.947 | 0.850 | | **<0.01** | **<0.01** |
| ***TEMRA*** | 0.783 | 0.708 | 0.809 | 0.793 | | **<0.01** |
| ***Treg*** | 0.771 | 0.759 | 0.694 | 0.788 | 0.778 | |

Table S2C.

| ***r/p value*** | ***CD8*** | ***CM*** | ***EM*** | ***Naïve*** | ***TEMRA*** |
|---|---|---|---|---|---|
| ***CD8*** | | **<0.01** | **<0.01** | **<0.01** | **<0.01** |
| ***CM*** | 0.928 | | **<0.01** | **<0.01** | **<0.01** |
| ***EM*** | 0.877 | 0.831 | | **<0.01** | **<0.01** |
| ***Naïve*** | 0.844 | 0.834 | 0.732 | | **<0.01** |
| ***TEMRA*** | 0.790 | 0.790 | 0.823 | 0.775 | |

Table S2D.

| ***r/p value*** | ***B cells*** | ***Memory*** | ***Naïve*** |
|---|---|---|---|
| ***B cells*** | | **<0.01** | **<0.01** |
| ***Memory*** | 0.511 | | **<0.01** |
| ***Naïve*** | 0.987 | 0.473 | |

Table S2E.

| ***r/p value*** | ***Monocytes*** | ***Classical*** | ***Intermediate*** | ***Non-classical*** |
|---|---|---|---|---|
| ***Monocytes*** | | **<0.01** | **<0.01** | **<0.01** |
| ***Classical*** | 0.915 | | **<0.01** | **<0.01** |
| ***Intermediate*** | 0.793 | 0.775 | | **<0.01** |
| ***Non-classical*** | 0.523 | 0.414 | 0.403 | |

Table S2F.

| ***r/p value*** | ***NK cells*** | ***CD56dimCD16+*** | ***CD56dimCD16-*** | ***CD56hiCD16+*** | ***CD56hiCD16-*** |
|---|---|---|---|---|---|
| ***NK cells*** | | **<0.01** | **<0.01** | **<0.01** | **<0.01** |
| ***CD56dimCD16+*** | 0.932 | | **<0.01** | **<0.01** | **<0.01** |
| ***CD56dimCD16-*** | 0.820 | 0.704 | | **<0.01** | **<0.01** |
| ***CD56hiCD16+*** | 0.739 | 0.636 | 0.704 | | **<0.01** |
| ***CD56hiCD16-*** | 0.835 | 0.700 | 0.743 | 0.814 | |

Table S2G.

| ***r/p value*** | ***Total*** | ***NK T*** | ***CM CD4*** | ***EM CD4*** | ***Naïve CD4*** | ***TEMRA CD4*** | ***Treg*** | ***CM CD8*** | ***EM CD8*** | ***Naïve CD8*** | ***TEMRA CD8*** | ***Memory B*** | ***Naïve B*** | ***CM*** | ***IM*** | ***NM*** | ***NK1*** | ***NK2*** | ***NK3*** | ***NK4*** |
|---|---|---|---|---|---|---|---|---|---|---|---|---|---|---|---|---|---|---|---|---|
| ***Total*** | | **<0.01** | **<0.01** | **0.013** | **<0.01** | **0.018** | **<0.01** | **<0.01** | **<0.01** | **<0.01** | **<0.01** | **<0.01** | **<0.01** | **<0.01** | **0.012** | **<0.01** | **<0.01** | **<0.01** | **0.002** | **<0.01** |
| ***NK T*** | 0.487 | | | **<0.01** | **<0.01** | **0.011** | **<0.01** | **<0.01** | **<0.01** | **<0.01** | **<0.01** | **<0.01** | **<0.01** | **<0.01** | **0.029** | **0.020** | **0.003** | **0.002** | **0.002** | **<0.01** |
| ***CM CD4*** | 0.576 | 0.579 | | **<0.01** | **<0.01** | **<0.01** | **<0.01** | **<0.01** | **<0.01** | **<0.01** | **<0.01** | **<0.01** | **0.022** | **<0.01** | **<0.01** | **0.040** | **0.005** | **<0.01** | **<0.01** | **<0.01** |
| ***EM CD4*** | 0.376 | 0.479 | 0.871 | | **<0.01** | **<0.01** | **<0.01** | **<0.01** | **<0.01** | **<0.01** | **<0.01** | **<0.01** | 0.189 | **<0.01** | **0.007** | 0.347 | **0.040** | 0.003 | **<0.01** | **<0.01** |
| ***Naïve CD4*** | 0.602 | 0.604 | 0.947 | 0.850 | | **<0.01** | **<0.01** | **<0.01** | **<0.01** | **<0.01** | **<0.01** | **<0.01** | **<0.01** | **<0.01** | **<0.01** | **0.032** | **0.001** | **<0.01** | **<0.01** | **<0.01** |
| ***TEMRA CD4*** | 0.358 | 0.383 | 0.708 | 0.809 | 0.793 | | **<0.01** | **<0.01** | **<0.01** | **<0.01** | **<0.01** | **<0.01** | 0.102 | **<0.01** | **0.008** | 0.256 | **0.011** | **<0.01** | **<0.01** | **<0.01** |
| ***Treg*** | 0.452 | 0.486 | 0.759 | 0.694 | 0.788 | 0.778 | | **<0.01** | **<0.01** | **<0.01** | **<0.01** | **<0.01** | **0.012** | **<0.01** | **0.014** | 0.289 | **0.002** | **<0.01** | **<0.01** | **<0.01** |
| ***CM CD8*** | 0.512 | 0.569 | 0.890 | 0.775 | 0.866 | 0.695 | 0.736 | | **<0.01** | **<0.01** | **<0.01** | **<0.01** | **0.009** | **<0.01** | **<0.01** | 0.091 | **0.004** | **<0.01** | **<0.01** | **<0.01** |
| ***EM CD8*** | 0.466 | 0.494 | 0.813 | 0.872 | 0.805 | 0.824 | 0.649 | 0.831 | | **<0.01** | **<0.01** | **<0.01** | **0.035** | **<0.01** | **<0.01** | 0.156 | **0.002** | **<0.01** | **<0.01** | **<0.01** |
| ***Naïve CD8*** | 0.531 | 0.510 | 0.864 | 0.800 | 0.875 | 0.730 | 0.798 | 0.834 | 0.732 | | **<0.01** | **<0.01** | **0.023** | **<0.01** | **0.005** | **0.019** | **0.004** | **<0.01** | **<0.01** | **<0.01** |
| ***TEMRA CD8*** | 0.471 | 0.466 | 0.741 | 0.737 | 0.828 | 0.883 | 0.859 | 0.790 | 0.823 | 0.775 | | **<0.01** | **<0.01** | **<0.01** | **<0.01** | 0.324 | **<0.01** | **<0.01** | **<0.01** | **<0.01** |
| ***Memory B*** | 0.546 | 0.494 | 0.651 | 0.583 | 0.719 | 0.704 | 0.695 | 0.666 | 0.735 | 0.674 | 0.783 | | **<0.01** | **<0.01** | **<0.01** | **0.004** | **<0.01** | **<0.01** | **<0.01** | **<0.01** |
| ***Naïve B*** | 0.480 | 0.436 | 0.349 | 0.204 | 0.439 | 0.253 | 0.379 | 0.393 | 0.323 | 0.346 | 0.456 | 0.473 | | 0.010 | 0.314 | 0.077 | **0.018** | 0.003 | 0.024 | 0.017 |
| ***CM*** | 0.624 | 0.511 | 0.742 | 0.627 | 0.759 | 0.512 | 0.630 | 0.653 | 0.674 | 0.688 | 0.646 | 0.716 | 0.387 | | **<0.01** | **0.006** | **<0.01** | **<0.01** | **<0.01** | **<0.01** |
| ***IM*** | 0.378 | 0.333 | 0.458 | 0.408 | 0.509 | 0.401 | 0.374 | 0.432 | 0.527 | 0.420 | 0.503 | 0.647 | 0.157 | 0.775 | | **0.007** | **<0.01** | **0.002** | **0.002** | **<0.01** |
| ***NM*** | 0.635 | 0.353 | 0.314 | 0.147 | 0.327 | 0.177 | 0.166 | 0.261 | 0.220 | 0.356 | 0.154 | 0.435 | 0.273 | 0.414 | 0.403 | | **0.005** | **0.023** | 0.115 | 0.062 |
| ***NK1*** | 0.604 | 0.447 | 0.419 | 0.314 | 0.472 | 0.385 | 0.463 | 0.433 | 0.465 | 0.432 | 0.532 | 0.610 | 0.359 | 0.611 | 0.531 | 0.423 | | **<0.01** | **<0.01** | **<0.01** |
| ***NK2*** | 0.681 | 0.453 | 0.571 | 0.445 | 0.682 | 0.556 | 0.648 | 0.601 | 0.567 | 0.584 | 0.731 | 0.703 | 0.449 | 0.686 | 0.462 | 0.346 | 0.704 | | **<0.01** | **<0.01** |
| ***NK3*** | 0.461 | 0.468 | 0.613 | 0.549 | 0.656 | 0.643 | 0.676 | 0.634 | 0.628 | 0.584 | 0.706 | 0.611 | 0.343 | 0.581 | 0.464 | 0.244 | 0.636 | 0.704 | | **<0.01** |
| ***NK4*** | 0.588 | 0.556 | 0.797 | 0.699 | 0.793 | 0.629 | 0.740 | 0.740 | 0.732 | 0.739 | 0.743 | 0.682 | 0.363 | 0.836 | 0.593 | 0.287 | 0.700 | 0.743 | 0.814 | |

Table S2H.

***Table S2.*** Correlation of total LC3B-II flux with major populations and subpopulations analyzed by correlation matrix with data presented as Spearman r and p value as follows: Total with major populations (A), T cells (B), CD4 T cells (C), CD8 T cells (D), B cells (E), monocytes (F), NK cells (G), total LC3B-II flux with subpopulations (H). CM (classical monocytes), IM (intermediate monocytes), NM (non-classical monocytes), NK1 ($CD56^{dim}CD16^{+}$ NK cells), NK2 ($CD56^{dim}CD16^{-}$ NK cells), NK3 ($CD56^{hi}CD16^{+}$NK cells), NK4 ($CD56^{hi}CD16^{-}$ NK cells).

| LC3BII flux | | | Total (n=43) | Males (n=19) | Female (n=24) | p value |
|---|---|---|---|---|---|---|
| ***Total*** | | | 41.4 (0.1-1427) | 24.9(0.1-742) | 61.5(0.1-1427) | 0.059 |
| ***T cells*** | | | 15.8 (0-96.65) | 8.4 (0-70) | 21.75 (0-96.65) | 0.059 |
| | *NKT cells* | | 34 (0-571) | 12.8 (0-539) | 40.75 (0-571) | 0.16 |
| | *CD4 T cells* | | 13.9 (0-100.6) | 4.5 (0-61) | 18 (0-100.6) | 0.14 |
| | | Central memory | 14.9 (0-93.2) | 9.4 (0-63) | 17.35 (0-93.2) | 0.14 |
| | | Effector memory | 14 (0-84) | 7.5 (0-79.7) | 16.2 (0-84) | 0.49 |
| | | Naïve | 14.5 (0-113.3) | 7.3 (0-64) | 15.2 (0-113.3) | 0.21 |
| | | TEMRA | 15.11 (0-106) | 13.1 (0-80.8) | 15.56 (0-106) | 0.79 |
| | | Regulatory | 9.3 (0-177.72) | 7.1 (0-56.8) | 12.98 (0-177.72) | 0.49 |
| | *CD8 T cells* | | 15 (0-79.2) | 86. (0-66.8) | 19 (0-79.2) | 0.15 |
| | | Central memory | 14.7 (0-407) | 12.8 (0-61.4) | 16.7 (0-407) | 0.19 |
| | | Effector memory | 11 (0-82.1) | 6.5 (0-75) | 14.73 (0-82.1) | 0.3 |
| | | Naïve | 9.6 (0-135.1) | 3 (0-74) | 15.5 (0-135.1) | 0.057 |
| | | TEMRA | 15 (0-89.4) | 12.6 (0-74.1) | 15.45 (0-89.4) | 0.9 |
| ***B cells*** | | | 41.1 (0-120.1) | 46.9 (0-110) | 32.75 (0-120.1) | 0.98 |
| | *Naïve* | | 36 (0-148) | 47.5 (0-134) | 28.55 (0-148) | 0.79 |
| | *Memory* | | 18.62 (0-126.9) | 7 (0-87.2) | 20.96 (0-126.9) | 0.17 |
| ***Monocytes*** | | | 10.9 (0-339) | 3 (0-93) | 18.39 (0-339) | **0.03** |
| | *Classical monocytes* | | 10.2 (0.79-21.8) | 8.41 (0.79-17.3) | 10.35 (1.67-21.8) | 0.62 |
| | *Intermediate monocytes* | | 21 (0-259) | 21 (0-111) | 20.195 (0-259) | 0.36 |
| | *Non-classical monocytes* | | 78 (0-3175) | 45 (0-174) | 180 (0-3175) | **<0.01** |
| ***NK cells*** | | | 18.49 (0-711) | 11.9 (0-198) | 22.31 (0-711) | 0.23 |
| | $CD56^{dim}$ $CD16^{+}$ | | 23.65 (0-1034) | 24.3 (0-257) | 17.86 (0-1034) | 0.46 |
| | $CD56^{dim}$ $CD16^{-}$ | | 15 (0-234) | 8.8 (0-132.5) | 17.15 (0-234) | 0.21 |
| | $CD56^{hi}$ $CD16^{+}$ | | 15.6 (0-217) | 15.6 (0-216) | 15.57 (0-217) | 0.47 |
| | $CD56^{hi}$ $CD16^{-}$ | | 9.38 (0-67) | 7 (0-64) | 10.79 (0-67) | 0.38 |

***Table S3.*** Sex differences for autophagic flux in different cell populations analyzed by Mann-Whitney test.

| Reagents or Resources | Source | Identifier |
| --- | --- | --- |
| ***Antibodies*** | | |
| BD Horizon BB700 Mouse Anti-Human CD127 | BD Biosciences | 566398<br>Clone HIL-7R-M21 |
| BD Horizon BUV395 Mouse Anti-Human CD4 | BD Biosciences | 563550<br>Clone K3 |
| BD Horizon BUV805 Mouse Anti-Human CD27 | BD Biosciences | 569167<br>Clone L128 |
| BD Horizon BV421 Mouse Anti-Human CD45 | BD Biosciences | 563879<br>Clone HI30 |
| BD Horizon BV480 Rat Anti-Human CCR7 (CD197) | BD Biosciences | 566099<br>Clone 3D12 |
| BD Horizon BV786 Mouse Anti-Human CD8 | BD Biosciences | 563823<br>Clone RPA-T8 |
| BD Horizon Fixable Viability Stain780 | BD Biosciences | 565388 |
| BD Horizon PE-CF594 Mouse Anti-Human CD19 | BD Biosciences | 562294<br>Clone HIB19 |
| BD OptiBuild BUV496 Mouse Anti-Human CD45RA | BD Biosciences | 741182<br>Clone 5H9 |
| BD OptiBuild BUV615 Mouse Anti-Human CD16 | BD Biosciences | 751323<br>Clone B73.1 |
| BD OptiBuild BV750 Mouse Anti-Human CD25 | BD Biosciences | 747290<br>Clone 2A3 |
| BD Pharmingen FITC Mouse Anti-Human CD3 | BD Biosciences | 555339<br>Clone HIT3a |
| BD Pharmingen FITC Mouse Anti-Human CD56 | BD Biosciences | 555516<br>Clone B159 |
| BD Pharmingen PE-Cy7 Mouse Anti-Human CD14 | BD Biosciences | 557742<br>Clone M5E2 |
| LC3B (E5Q2K) Mouse mAb (Alexa Fluor® 647 Conjugate) | Cell Signaling Technology | 18577S |
| ***Chemicals and consumables*** | | |
| Bafilomycin A1 | Selleck Chemicals | S1413 |
| BD Horizon Brilliant Stain Buffer Plus | BD Biosciences | 566385 |
| BD™ Cytometer Setup and Tracking Beads | BD Biosciences | 642412 |
| Bovine serum albumin | Sigma Aldrich | A9647-100G |
| Chloroquine diphosphate | Sigma Aldrich | C6628 |
| Dulbecco's modifed Eagle's medium (DMEM, high glucose, no glutamine) | Thermo Fisher Scientific | 11960044 |
| Dulbecco's phosphate-buffered saline (DPBS) | GIBCO, Thermo Fisher Scientific | 14 190 136 |
| Fetal bovine serum (undialyzed) | Life Technologies | 10 099–141 |
| Fetal bovine serum, dialyzed (d.FBS) | Thermo Fisher Scientific | A3382001 |
| Lithium heparin Vacu3tte tube 9mL | Greiner Bio-One | 455.084 |
| Lymphoprep | StemCell Technologies | 07811 |
| Neutral formalin | Thermo Fisher Scientific | BSPFS426.2.5 |
| Red blood cell lysis buffer | BD Biosciences | 555 899 |

| | | |
|---|---|---|
| RPMI 1640 medium | Life Technologies | R8758 |
| RPMI 1640 medium w/o amino acids, sodium phosphate (powder) | US Biological | R8999-04A |
| RPMI 1640 medium, no glucose | Thermo Fisher Scientific | 11879020 |
| Saponin from quillaja bark | Sigma Aldrich | S4521-10G |
| Glucose solution | Thermo Fisher Scientific | A2494001 |
| Sodium bicarbonate | Sigma Aldrich | S5761 |
| Sodium phosphate dibasic anhydrous | Sigma Aldrich | S9763 |
| L-Aspartic acid | Sigma Aldrich | A9256-100G |
| L-Serine | Sigma Aldrich | S4311-25G |
| L-Threonine | Sigma Aldrich | T8625-10G |
| L-Asparagine | Sigma Aldrich | A0884-25G |
| L-Cysteine | Sigma Aldrich | C7352-25G |
| L-Alanine | Sigma Aldrich | A7469-25G |
| L-Methionine | Sigma Aldrich | M9625-25G |
| L-Phenylalanine | Sigma Aldrich | P2126-100G |
| L-Tyrosine | Sigma Aldrich | T3754-50G |
| L-Tryptophan | Sigma Aldrich | T0254-25G |
| L-Leucine | Sigma Aldrich | L8912 |
| L-Valine | Sigma Aldrich | V0513 |
| L-Isoleucine | Sigma Aldrich | I7403 |
| L-Lysine | Sigma Aldrich | L5501 |
| L-Histidine | Sigma Aldrich | H8000 |
| L-Arginine | Sigma Aldrich | A8094 |
| L-Glutamic acid (monosodium salt hydrate) | Sigma Aldrich | G5889 |
| L-Proline | Sigma Aldrich | P03080 |
| Glycine | Millipore | VP709001610 |
| L-Glutamine | Sigma Aldrich | G7513 |
| ***Software and Algorithms*** | | |
| Flowjo 10.8.0 for Window | Tree Star | |
| GRAPHPAD PRISM, version 10.1.0 for Windows | GraphPad | |
| R version 4.3.2 | The R Foundation www.r-project.org | |
| RStudio | www.rstudio.com | |

***Table S4***. Reagents used in the study.

| Amino Acids | Molecular Weight | Concentration (mM) |
| --- | --- | --- |
| Glycine | 75 | 0.133 |
| L-Arginine | 174 | 1.149 |
| L-Asparagine | 132 | 0.379 |
| L-Aspartic acid | 133 | 0.15 |
| L-Cystine 2HCl | 313 | 0.208 |
| L-Glutamic Acid | 147 | 0.136 |
| L-Glutamine* | 146 | 2.055 |
| L-Histidine | 155 | 0.097 |
| L-Isoleucine | 131 | 0.382 |
| L-Leucine | 131 | 0.382 |
| L-Lysine hydrochloride | 183 | 0.219 |
| L-Methionine | 149 | 0.101 |
| L-Phenylalanine | 165 | 0.091 |
| L-Proline | 115 | 0.174 |
| L-Serine | 105 | 0.286 |
| L-Threonine | 119 | 0.168 |
| L-Tryptophan | 204 | 0.025 |
| L-Tyrosine disodium salt dihydrate | 261 | 0.111 |
| L-Valine | 117 | 0.171 |
| L-Alanine | 89.09 | 0.4 |

***Table S5***. Amino acids and concentrations spiked into $aa^{+}$ condition in Fig. 5
